# Origin of the mobile di-hydro-pteroate synthase gene determining sulfonamide resistance in clinical isolates

**DOI:** 10.1101/472472

**Authors:** Miquel Sánchez-Osuna, Pilar Cortés, Jordi Barbé, Ivan Erill

## Abstract

Sulfonamides are synthetic chemotherapeutic agents that work as competitive inhibitors of the di-hydro-pteroate synthase (DHPS) enzyme, encoded by the *folP* gene. Resistance to sulfonamides is widespread in the clinical setting and predominantly mediated by plasmid- and integron-borne *sul1-3* genes encoding mutant DHPS enzymes that do not bind sulfonamides. In spite of their clinical importance, the genetic origin of *sul1-3* genes remains unknown. Here we analyze *sul* genes and their genetic neighborhoods to uncover *sul* signature elements that enable the elucidation of their genetic origin. We identify a protein sequence Sul motif associated with sul-encoded proteins, as well as consistent association of a phosphoglucosamine mutase gene (*glmM*) with the *sul2* gene. We identify chromosomal *folP* genes bearing these genetic markers in two bacterial families: the *Rhodobiaceae* and the *Leptospiraceae*. Bayesian phylogenetic inference of FolP/Sul and GlmM protein sequences clearly establishes that *sul1-2* and *sul3* genes originated as a mobilization of *folP* genes present in, respectively, the *Rhodobiaceae* and the *Leptospiraceae*, and indicate that the *Rhodobiaceae folP* gene was transferred from the *Leptospiraceae*. Analysis of %GC content in *folP*/*sul* gene sequences supports the phylogenetic inference results and indicates that the emergence of the Sul motif in chromosomally-encoded FolP proteins is ancient and considerably predates the clinical introduction of sulfonamides. *In vitro* assays reveal that both the *Rhodobiaceae* and the *Leptospiraceae*, but not other related chromosomally-encoded FolP proteins confer resistance in a sulfonamide-sensitive *Escherichia coli* background, indicating that the Sul motif is associated with sulfonamide resistance. Given the absence of any known natural sulfonamides targeting DHPS, these results provide a novel perspective on the emergence of resistance to synthetic chemotherapeutic agents, whereby preexisting resistant variants in the vast bacterial pangenome may be rapidly selected for and mobilized upon the clinical introduction of novel chemotherapeuticals.

## 1 Introduction

Antibiotic resistance is a pressing problem in modern healthcare [1,2]. Bacterial cells present several mechanisms to cope with exposure to antibiotics or chemotherapeutic agents, which may be acquired through mutation or, most frequently, via lateral gene transfer on mobile genetic elements [3]. These mechanisms include modification of the antimicrobial target, degradation or chemical modification of the antimicrobial molecule, targeted reduction of antimicrobial uptake, active export of the antimicrobial through efflux pumps and use of alternate pathways and enzymes [3].

It is widely accepted that many antibiotic resistance genes present today in pathogenic bacteria originated from homologs evolved over eons in either the microbes that naturally produce the antibiotics or their natural competitors [4]. When coupled with the high plasticity of bacterial genomes and their co-existence with a large variety of genetic mobile elements, the availability of a readily evolved pool of antibiotic resistance genes set the stage for the rapid proliferation of multi-resistant strains in the clinical setting shortly after the commercial introduction of antibiotics [4]. In contrast, the origins of resistance against chemotherapeutic agents are harder to pinpoint. Since these were designed *in vitro*, it seems unlikely that a large pool of genes conferring resistance to chemotherapeutic agents existed before their introduction. After their discovery in the 1960’s, resistance to quinolones was initially rare and limited to chromosomal mutations in DNA gyrase, topoisomerase IV or efflux pumps [5]. However, in the early 2000’s plasmid-borne *qnr* genes were first detected and spread rapidly to clinical pathogens. Qnr is a member of the pentapeptide repeat family and was shown to confer resistance by binding to DNA gyrase and limiting the effect of quinolone drugs. The origin of plasmid-borne *qnr* genes has been traced to environmental homologs and these are thought to have derived from genes originally targeting antibiotics, such as microcin B17 [6].

Aryl sulfonamides are synthetic antibacterial compounds presenting a similar structure to para-amino benzoic acid (PABA), and containing a sulfonamide group linked to an aromatic group. Commonly referred to as sulfonamides or sulfa drugs due to their clinical relevance, synthetic aryl sulfonamides function as competitive inhibitors of the di-hydro-pteroate synthase (DHPS) enzyme, encoded in bacteria by the *folP* gene [7]. DHPS participates in folate synthesis using PABA as a substrate, and the competitive inhibition of DHPS by sulfonamides results in growth arrest [7,8]. Experiments in mice in the 1930’s demonstrated the effectiveness of sulfonamide against bacteria, and sulfonamide became the first antibacterial chemotherapeutic to be used systemically [9,10]. It remained in use throughout World War II, but by the end of the 1940’s resistant strains started to emerge and sulfonamides were rapidly displaced in favor of the newly discovered antibiotics [7,11].

Resistance to sulfonamide through increased production of PABA was reported in the early 1940’s [12], but the most commonly reported mechanism of sulfonamide resistance are mutations to the chromosomal *folP* gene [7,13]. Mutations to the chromosomal *folP* gene have been shown to provide varying degrees of trade-off between resistance and efficient folate synthesis, decreasing DHPS affinity for sulfonamide while maintaining or increasing its affinity for PABA [7]. These mutations have occurred independently in multiple bacterial genera and target multiple conserved areas of the DHPS protein [7]. However, similar mutational profiles, such as two-amino acid insertions in *Neisseria meningitidis* and *Streptococcus pneumoniae*, have been reported [14,15], and in both these genera there is evidence of extensive recombination within *folP* genes [16,17].

In spite of the multiple instances of chromosomal *folP* resistant variants, clinical resistance to sulfonamides is predominantly plasmid-borne and mediated by *sul* genes encoding alternative sulfonamide-resistant DHPS enzymes [7]. Four different *sul* genes have been described to date, with *sul1* and *sul2* being the predominant forms in clinical isolates [18]. The *sul1* gene is typically found in class 1 integrons and linked to other resistance genes [18], whereas *sul2* is usually associated to non-conjugative plasmids of the IncQ group [19] and to large transmissible plasmids like pBP1 [20]. The *sul3* gene was characterized in the *Escherichia coli* conjugative plasmid pVP440. It was shown to be flanked by two copies of the insertion element IS15Δ/26 and to be widespread in *E. coli* isolates from pigs in Switzerland [21]. Recently, a *sul4* gene was identified in a systematic prospection of class 1 integron-borne genes in Indian river sediments, but this *sul* variant has not yet been detected in clinical isolates. Genomic context analyses revealed that the *sul4* gene had been recently mobilized and phylogenetic inference pinpointed its putative origin as part of the folate synthesis cluster in the Chloroflexi phylum [22].

Despite the importance of sulfonamides in human and animal therapy, the putative origin of the three *sul* genes that account for the vast majority of reported clinical resistance to sulfonamide remains to be elucidated. In this work we leverage comparative genomics, phylogenetic analysis and *in vitro* determination of minimal inhibitory concentrations (MIC) of sulfamethoxazole to unravel the origin of the *sul1, sul2* and *sul3* genes. Our analysis indicates that chromosomally-encoded *folP* genes conferring resistance to sulfonamide originated in members of the *Leptospiraceae* family and were transferred to the Alphaproteobacteria *Rhodobiaceae* family more than 500 million years ago. These isolated sources of chromosomally-encoded sulfonamide-resistant DHPS were mobilized independently following the commercial introduction of sulfonamides, leading to the broadly disseminated sul1, *sul2* and *sul3* resistance genes. Our results hence indicate that resistance to synthetic chemotherapeutic agents may be available in the form of chromosomally-encoded variants among the extremely diverse bacterial domain, and can be rapidly disseminated upon the release of novel synthetic drugs.

## 2 Materials and methods

### 2.1 Data collection

FolP, GlmM and Sul1-3 homologs were identified in complete GenBank sequences through BLASTP [23] using the *E. coli* FolP (WP_000764731) and GlmM (WP_000071134) proteins as the query. Putative homologs were detected as BLASTP hits passing stringent e-value (<1e-20) and query coverage (75%) thresholds. FolP and GlmM chromosomally-encoded proteins were identified on a representative genome of all bacterial orders with complete genome assemblies on RefSeq, of each bacterial family for the Proteobacteria, of any bacterial species where chromosomally-encoded sulfonamide resistance mutants had been reported, and on all available complete genomes for clades of interest (*Rhodobiaceae*, Spirochaetes and Chlamydiae) (Supplementary material 1). All protein coding gene sequences for these genomes were downloaded for %GC analysis. Sul proteins encoded by mobile *sul* genes were identified on complete plasmid, transposon and integron GenBank sequences.

### 2.2 Identification and visualization of Sul-like signatures in FolP sequences

To identify sequence motifs associated with Sul proteins, we performed a CLUSTALW alignment using a non-redundant (<99% identity) subset of the Sul1-3 homologous sequences detected previously and FolP sequence sampled from each bacterial clade. Following visual inspection of the resulting alignment, a Sul-like motif conserved in several chromosomally-encoded FolP proteins was visualized using iceLogo [24] and a consensus motif was derived and encoded into a PROSITE-format pattern. The inferred PROSITE pattern was used to seed a Pattern Hit Initiated BLAST search against the NCBI non-redundant Protein database using as a query the protein sequences of Sul1-3 reported in the literature (WP_001336346, WP_010890159, WP_000034420) and conservative e-value (<1e-20) and query coverage (75%) limits. Only chromosomal hits with the identified signature characteristic of *sul* gene products were retained for further analysis.

### 2.3 Multiple sequence alignment and phylogenetic inference

For phylogenetic inference, multiple sequence alignments of identified FolP/Sul1-3 and GlmM homologous sequences were performed with CLUSTALW [25] using variable (5, 10 and 25) gap opening penalties. These alignments were then integrated with local LALIGN alignments with T-COFFEE [26], and the resulting alignment was trimmed using the “less stringent selection” parameters of the Gblocks online service [27,28]. Bayesian phylogenetic inference on trimmed alignments was performed with MrBayes [29]. Four Metropolis-Coupled Markov Chain Monte Carlo runs with four independent chains were carried out for 30,000,000 generations, and the resulting consensus tree was plotted with FigTree.

### 2.4 DNA sequence analyses

Analysis of %GC in synonymous and non-synonymous patterns and *K_a_*/*K_s_* divergence were performed according to the Nei-Gojobori computation method [30] and the standalone PAL2NAL program for codon-based alignments [31], using custom Python scripts for pipelining. Analyses of %GC content were performed on all sampled bacterial genomes, computing genome-wide %GC statistics and comparing them to *folP* estimates. Analyses of *K_a_*/*K_s_* divergence were performed on pair-wise alignments of the N- and C-terminal ends of the *glmM* gene sequence of all sampled bacterial groups. One-sided Mann-Whitney U-tests were performed using GraphPad Prism to determine whether differences between *folP* and chromosomal %GC content were different in the presence and absence of Sul-like signature motifs, and whether the N- and C-terminal regions presented different mutational profiles. The scripts used for the analysis are available at the GitHub ErillLab repository. Amelioration times were estimated using the Ameliorator program [32] under different selection modes. *K_a_* and *K_s_* values were estimated from pairwise alignments of orthologs between the *Parvibaculum lavamentivorans* and *Leptospira interrogans* genomes as determined by the OMA Orthology database [33] and species divergence times were inferred from published molecular clock phylogenies [34].

### 2.5 Cloning, transformation and complementation of the *folP* gene for broth microdilution assays

The *L. interrogans* serovar Lai str. 56601 *folP* and *Chlamydia trachomatis* D/UW-3/CX *folKP* gene were synthesized and adapted to E. *coli* codon usage at ATG:biosynthetics GmbH, Germany; whereas *P. lavamentivorans* DS-1 (DSMZ 13023) and *Rhodobacter sphaeroides* 2.4.1 (gently provided by Professor S. Kaplan; Health Science Center. University of Texas) *folP* genes were amplified from genomic DNA. The *sul2* gene was amplified from the RSF1010 plasmid (Josep Antón, Instituto de Biotecnología y Biomedicina) [35,36] and used as a positive control. The *folP/sul* genes were subcloned into the expression vector pUA1108 using NdeI and BamHI (Supplementary material 2), as previously described [37] and the recombinant plasmids were then transformed into competent E. *coli* K-12 (CGSC 5073). Antimicrobial susceptibility testing of sulfamethoxazole (Sigma-Aldrich) for the strains containing the *folP/sul* genes was determined as described using broth microdilution tests in Mueller-Hinton broth (MH) with half serial dilutions of sulfamethoxazole ranging from 512 to 0.125 mg/L [38].

## 3 Results

### 3.1 Identification of putative chromosomal origins for *sul1-3* genes

To identify putative chromosomal homologs of *sul1-3* genes, we performed a multiple sequence alignment including any protein sequences with at most 99% similarity to those encoded by *sul1-3* genes reported in the literature and by chromosomal *folP* genes from a representative of each bacterial order. Inspection of the resulting alignment (Figure 1A; Supplementary material 3) revealed the presence of a two-amino acid insertion in proteins encoded by *sul1-3* genes that is not present in those encoded by *sul4* or the analyzed chromosomal *folP* genes. This two-amino acid insertion is located in a conserved region of the FolP protein (residues R171-N211 of the *E. coli* FolP protein [WP_000764731]) that presents other signature changes in sul-encoded proteins with respect to chromosomally-encoded FolP proteins (Figure 1AB; Supplementary material 3) [**39,40**]. We derived a PROSITE-format pattern (Supplementary material 4) of the identified Sul motif to seed a Pattern Hit Initiated BLAST search against the NCBI non-redundant (NR) protein database. This search identified several proteins encoded by *Rhodobiaceae* family members that presented a similar insertion pattern. BLASTP searches with these *Rhodobiaceae* FolP sequences matched proteins in several members of the *Leptospiraceae* and the Chlamydiae. However, analysis of the resulting multiple sequence alignment showed that only the *Leptospiraceae* FolP protein sequences displayed the identified two-amino acid insertion pattern (Supplementary material 5). Heretofore, we refer to these chromosomally-encoded FolP proteins containing the signature Sul motif as FolP*, and to their encoding gene as *folP*\*.

**Figure 1.**
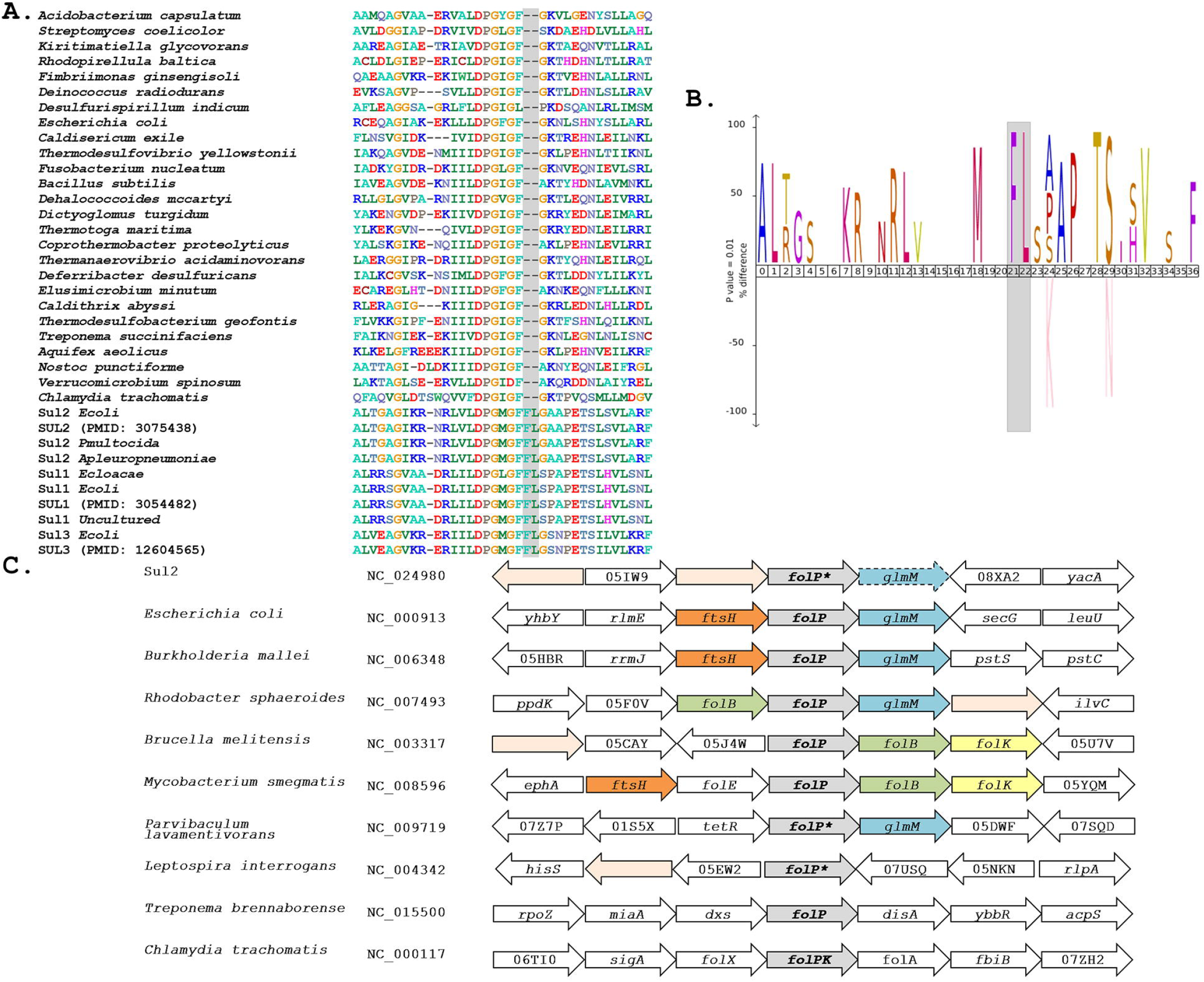
(A) Segment of the multiple sequence alignment including any *sul* genes with at most 90% similarity to reported *sul* genes and a representative chromosomal *folP* gene for all bacterial phyla with complete genomes available in NCBI RefSeq. (B) IceLOGO highlighting the difference in amino acid frequency at each position of the region of the *folP* protein sequence containing the identified insertion between the multiple sequence alignment of *sul* gene products and the chromosomally-encoded FolP proteins. The upper part of the iceLOGO plot shows residues overrepresented in the sul-encoded FolP proteins; the bottom part shows residues overrepresented in chromosomally-encoded FolP proteins for all bacterial phyla with complete genomes available. Only differences with significant z-score under a confidence interval of 0.01 are shown. (C) Schematic representation of the genetic environment of *sul2* genes, similar arrangements in chromosomally-encoded *folP* genes of the Gammaproteobacteria, Betaproteobacteria and Alphaproteobacteria, and arrangements in other major phyla. Arrow boxes indicate coding regions. When available, gene names or NOG identifiers are provided. Boxes for *folP* genes containing the two-amino acid insertion are designated as *folP*\*.

In order to gain further insight into the possible chromosomal origins of *sul* genes, we performed tBLASTX searches against the NCBI RefSeq Genome Database using the genetic surroundings (5,000 bp) of sul1, *sul2* and *sul3* genes with at most 90% similarity to those reported in the literature (Supplementary material 6). This search did not return consistent results for the *sul1* and *sul3* genetic surroundings, but it identified a conserved gene fragment encoding the N-terminal region of the phosphoglucosamine mutase GlmM protein downstream of *sul2* in multiple plasmids harboring this resistance gene. These *sul2*-associated GlmM sequences lack the entire GlmM C-terminal region, including three of its functional domains [41], and it can therefore be safely assumed that they are not functional as phosphoglucosamine mutases. This genetic arrangement has been reported previously as a feature of *sul2* isolates [42,43], and it is strongly conserved in the genomic surroundings of chromosomal *folP* genes in the Gammaproteobacteria, the Betaproteobacteria and several Alphaproteobacteria lineages (Figure 1C). Analysis of the *folP* genetic surroundings in complete genomes of the Spirochaetes and the Alphaproteobacteria shows clear differences between the genes coding for the identified *Rhodobiaceae* and *Leptospiraceae* FolP* proteins harboring the two-amino acid insertion pattern and those without it (Figure 1C). The *Leptospiraceae* show a conserved arrangement with *folP* flanked by a peptidoglycan-associated lipoprotein and a tetratricopeptide repeat-containing domain protein, whereas in most other Spirochaetes *folP*\* is flanked by a 1-deoxy-D-xylulose-5-phosphate synthase and a diadenylate cyclase. In contrast, the Alphaproteobacteria yield several distinct syntenic regions for *folP*. In the *Rhodobiaceae, folP* is flanked by genes coding for either a FtsH-family metallopeptidase or a TetR-family transcriptional repressor and the phosphoglucosamine mutase *glmM*. In the Rhodobacterales, *folP*\* is flanked by a dihydroneopterin aldolase and *glmM*, but in the *Rhizobiales* it is flanked by a Zn-dependent proteoase and the dihydroneopterin aldolase. This last arrangement, in which the dihydroneopterin aldolase is followed by a 2-amino-4-hydroxy-6-hydroxymethyldihydropteridine diphosphokinase is also part of the genetic surroundings of *folP* in most Actinobacteria (Figure 1C).

### 3.2 Phylogenetic analysis of *sul/folP* and *glmM* genes

The presence of a signature two-amino acid insertion characteristic of *sul* gene products in chromosomally-encoded FolP* proteins and the identification of a genetic environment for *sul2* genes that is conserved in multiple bacterial genomes suggested that it might be possible to pinpoint the evolutionary origin of *sul* genes. To further investigate this possibility, we performed a rigorous phylogenetic analysis of FolP/Sul protein sequences. We sampled a representative genome of all bacterial orders with complete genome assemblies, of each bacterial family for the Proteobacteria and all available complete genomes for clades of interest (*Rhodobiaceae*, Spirochaetes and Chlamydiae), and we identified chromosomally-encoded FolP homologs in each of these genomes using BLASTP with the E. *coli* FolP protein as a query. We used a distance tree generated with CLUSTALW to identify and discard a set of protein sequences from duplicated *folP* genes in the Actinobacteria (Supplementary material 7), and we performed multiple sequence alignment and Bayesian phylogenetic reconstruction of the remaining FolP/Sul sequences with T-COFFEE and MrBayes (Supplementary material 8).

The resulting tree (Figure 2) provides strong support for the hypothesis that *sul1-3* genes originated in the *Rhodobiaceae* and *Leptospiraceae* families. In particular, the topology inferred by MrBayes suggests that the *Leptospiraceae folP*\* gene gave rise to both *sul3* and the *folP*\* gene encountered in the *Rhodobiaceae*, most likely through a lateral gene transfer event in an ancestor of this Alphaproteobacteria family. According to the reconstructed FolP phylogeny, the *Rhodobiaceae folP*\* gene was subsequently mobilized as *sul2*, and later evolved into the integron-borne *sul1* gene [44]. The fact that the *Leptospiraceae* FolP* sequences branch independently of other Spirochaetes sequences and immediately after the Chlamydiae suggests that the *Leptospiraceae folP*\* gene might have originated as a result of lateral gene transfer event from the Chlamydiae, and that it subsequently incorporated the signature two-amino acid insert present in sul-encoded DHPS proteins.

**Figure 2.**
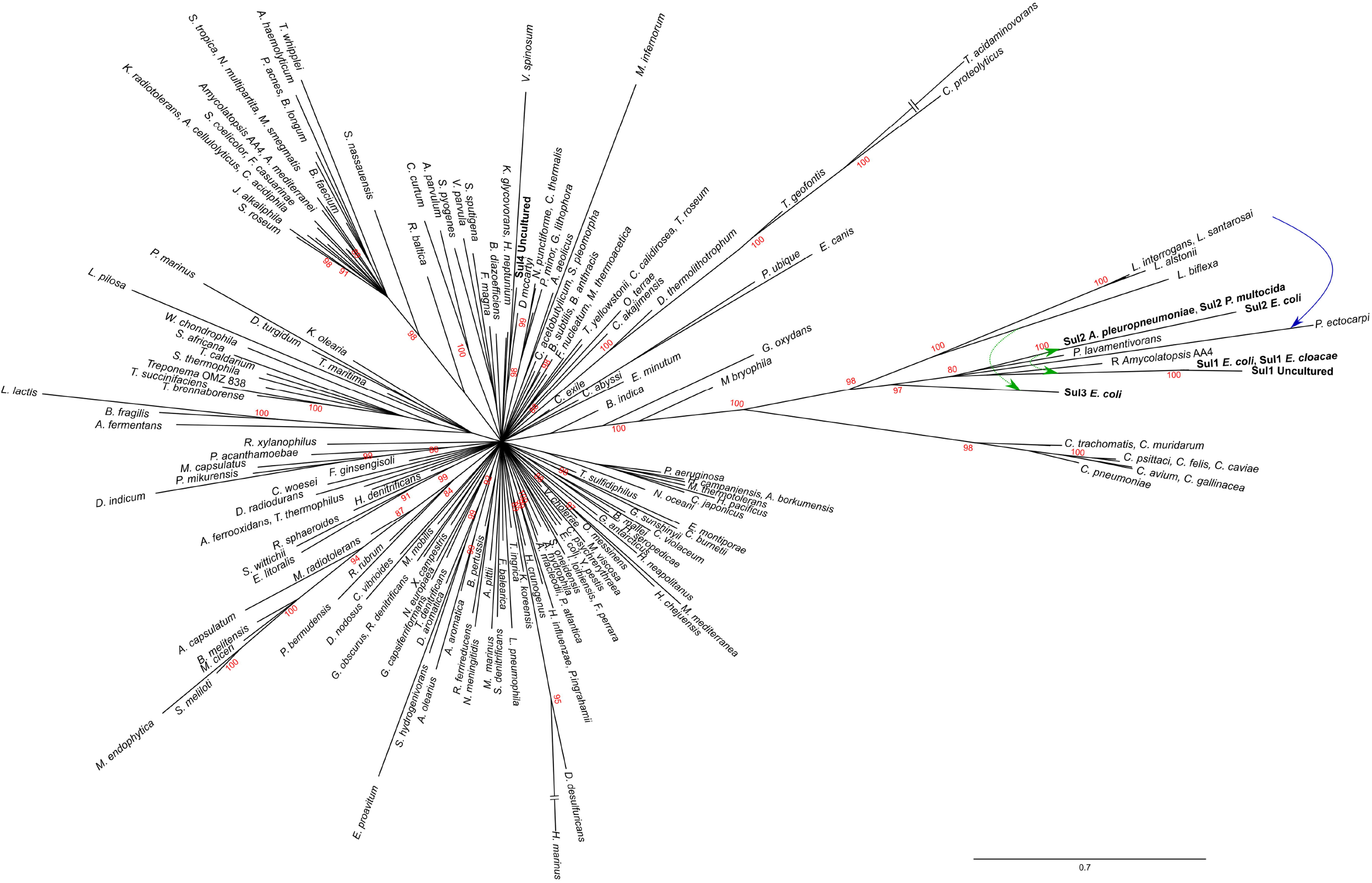
Consensus tree of Sul/FolP protein sequences. Branch support values are provided as Bayesian posterior probabilities. For clarity, only posterior probability values higher than 0.8 are displayed. Proposed lateral gene transfer and mobilization events are shown by means of superimposed continuous and dotted, respectively, arrows.

The existence of a genetic environment for *sul2* genes conserved in bacterial chromosomes provides the means to independently assess the likelihood of the evolutionary scenario inferred from the FolP phylogeny. Using the same sampling methods utilized for *sul/folP* protein products, we collected protein sequences for phosphoglucosamine mutase (GlmM) homologs and performed Bayesian phylogenetic inference on the aligned N-terminal regions. The resulting GlmM tree (Figure 3) provides further support for a *Rhodobiaceae* origin of the *sul2* gene, with the *sul2*-associated GlmM sequences branching with the *Rhodobiaceae* GlmM protein sequences deep within an otherwise monophyletic Alphaproteobacteria clade. Taken together, the consistent branching with the *Rhodobiaceae* of the protein sequences encoded by both *sul2* and its accompanying *glmM* gene fragment firmly establish this Alphaproteobacteria family as the chromosomal origin for the *sul2* gene. The phylogenetic evidence thus indicates that the *sul2* gene was excised with the N-terminal fragment of the *glmM* gene during the mobilization event that led to their incorporation into plasmid vectors. Given that the *folP-glmM* arrangement is only seen in the Proteobacteria, this also excludes the possibility that the *sul2* gene was mobilized directly from a *Leptospiraceae* background, where the *folP* gene presents an unrelated, yet conserved, genomic environment (Figure 1C).

**Figure 3.**
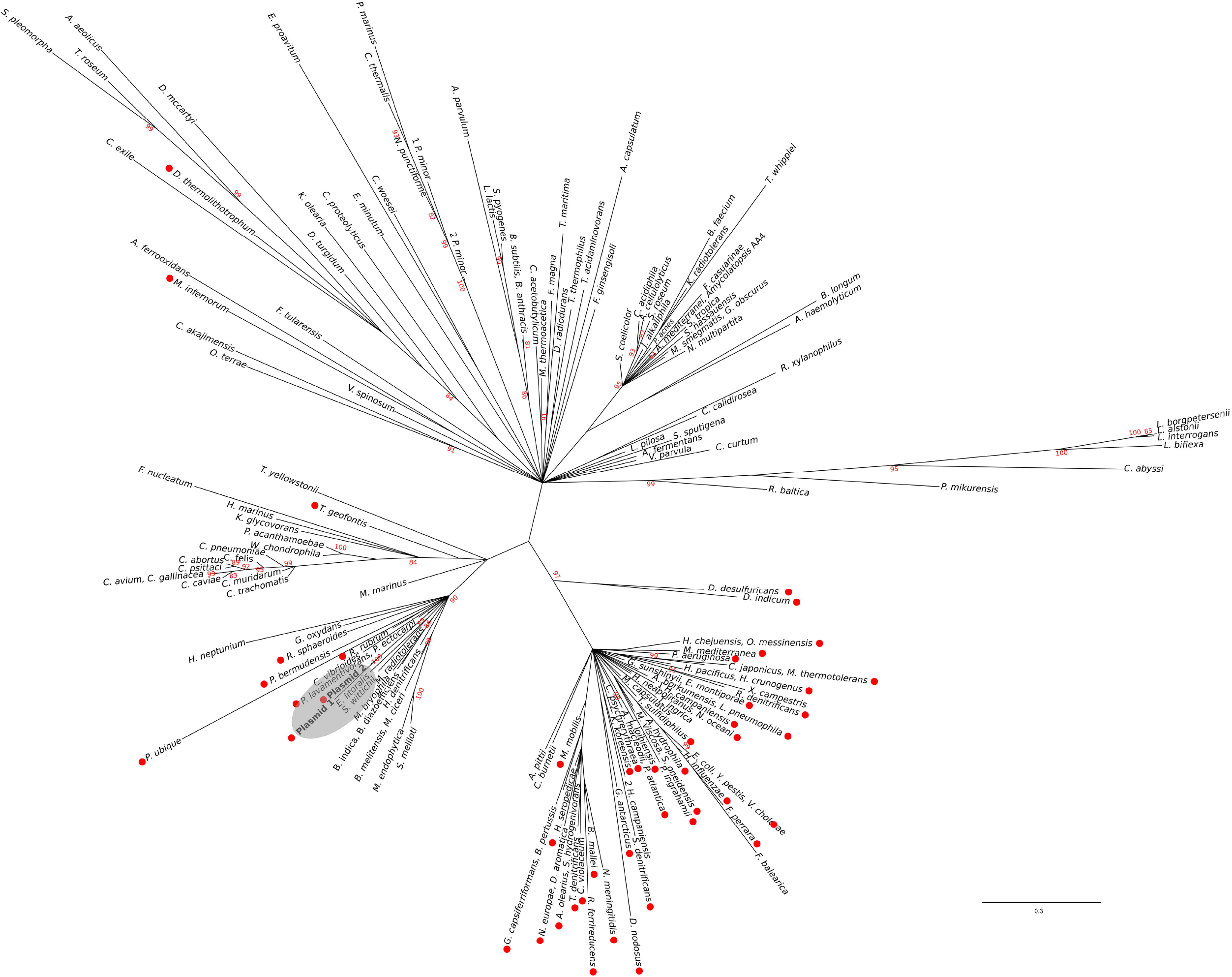
Consensus tree of N-terminal GlmM protein sequences. Branch support values are provided as Bayesian posterior probabilities. For clarity, only posterior probability values higher than 0.8 are displayed. The placement of *sul2*-encoded proteins is indicated by a shaded ellipse.

### 3.3 Analysis of *sul*/*folP* and *glmM* gene sequences

The phylogenetic analysis of FolP and GlmM sequences puts forward an evolutionary scenario wherein the *Leptospiraceae folP*\* was transferred to the members of the *Rhodobiaceae* family before being mobilized independently into the *sul3*- and sul1/2-harboring mobile genetic elements reported in sulfonamide-resistant clinical isolates. To further investigate this hypothesis, we undertook a systematic analysis of *folP* and *glmM* coding sequences. We compiled *folP* gene sequences for all the FolP proteins included in the phylogenetic analysis (Figure 2), as well as any *sul* gene sequences with less than 90% identity to those reported in the literature and any chromosomal *folP*\* genes encoding a DHPS with the signature Sul motif (Figure 1A) for which there were at least 1 Mbp of whole genome shotgun sequence data (Supplementary material 9). We computed the overall and codon-position %GC content on both the *folP*/*sul* coding sequences and all the available coding sequences in their respective genome assembly (Supplementary material 10). The %GC content data (Figure 4A) reveals that *sul1/2* sequences have a high %GC content (60.76 SD±1.42) that is consistent with their origin as mobilized *Rhodobiaceae folP*\* sequences (%GC content: 62.02 SD±2.22). Similarly, *sul3* sequences display a %GC content (38.14 SD±0.55) consistent with their mobilization from a *Leptospiraceae folP*\* background (39.39 SD±4.17). Together with the phylogenetic inference results, these data provide strong support for an independent mobilization of *sul1/2* and *sul3* genes from, respectively, *Rhodobiaceae* and *Leptospiraceae* chromosomal backgrounds.

**Figure 4.**
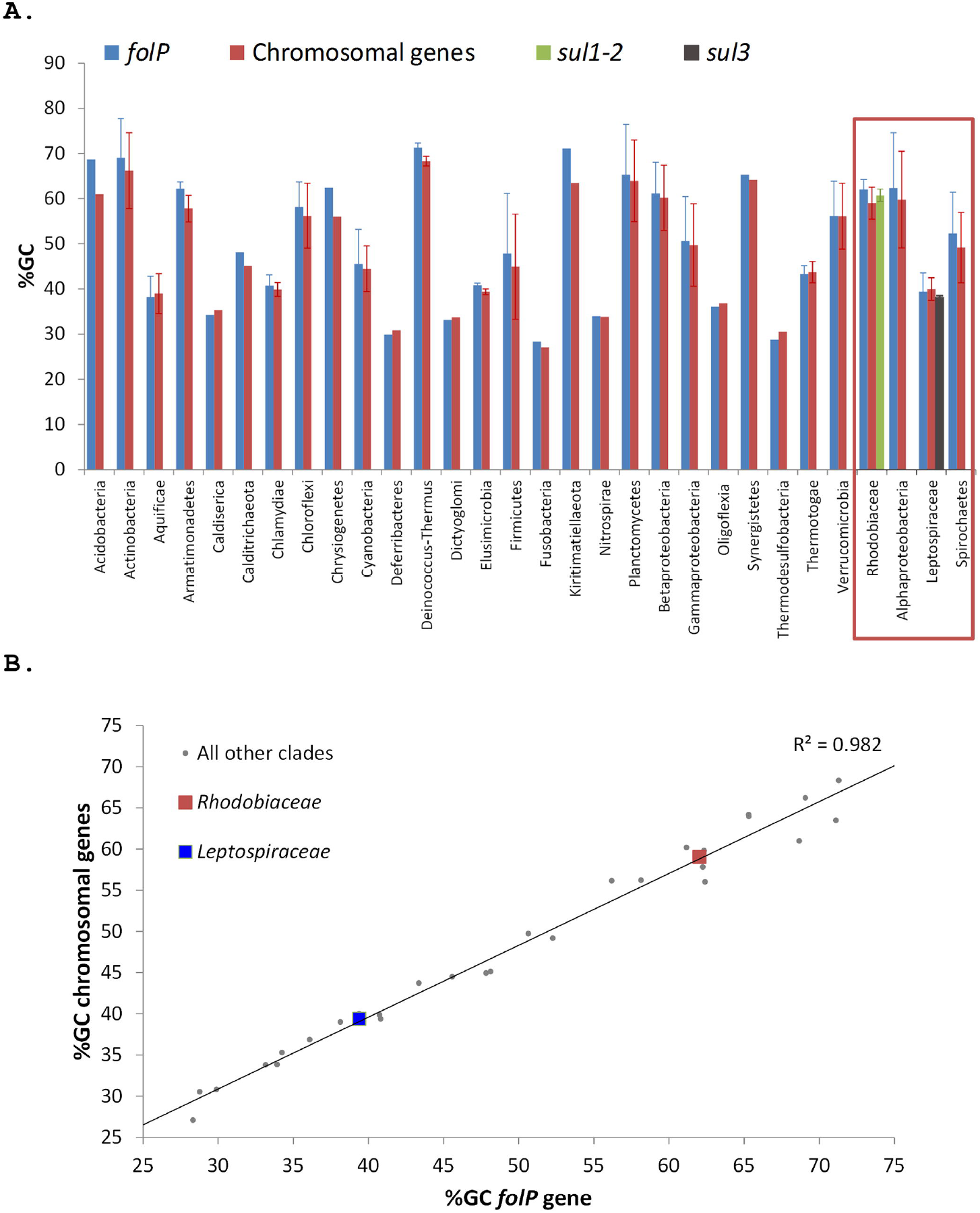
(A) %GC content of *folP* and all other chromosomal coding sequences in different clades. The %GC content of sul1/2 and sul3 genes is shown adjacent to that of the *Rhodobiaceae* and the *Leptospiraceae*. (B) Correlation between the %GC content of *folP* genes and that of all other coding sequences in their respective genomes. The data points corresponding to *folP*\* genes from the *Rhodobiaceae* and the *Leptospiraceae* are shown as squares.

The independent mobilization of *sul1/2* and *sul3* is underpinned by a preceding lateral gene transfer of *folP*\* from the *Leptospiraceae* into a *Rhodobiaceae* ancestor. In this context, the substantial divergence in %GC content between the chromosomal *folP*\* genes of both clades indicates a long process of amelioration. In fact, statistical analysis of the differences in codon position %GC content between *folP* genes and all available coding sequences in their respective genomes shows that *Leptospiraceae* and *Rhodobiaceae folP*\* genes encoding proteins with the Sul motif cannot be distinguished from other *folP* genes (one-sided Mann-Whitney U-test p > 0.05 for GC1, GC2 and GC3) (Figure 4B) (Supplementary material 10). We used Ameliorator [32] to estimate the time required for the observed amelioration via forward simulation from *Leptospiraceae* codon position %GC values. Even under assumptions of fast evolutionary change, the software provides a lower bound of 476 million years for the observed amelioration of the *Leptospiraceae folP*\* gene into the *Rhodobiaceae* one. Statistical analysis of synonymous and non-synonymous mutation patterns in the N- and C-terminal regions of the *glmM* gene also shows that mutation patterns in each region of the *Rhodobiaceae glmM* gene are indistinguishable from those observed in other *glmM* genes (one-sided Mann-Whitney U-test p > 0.05), indicating that the *glmM* gene fragment associated with *sul* genes was not transferred from a mobile element into the *Rhodobiaceae* (Supplementary material 11).

### 3.4 Sulfonamide resistance of chromosomal *folP* genes

Phylogenetic and sequence analysis results indicate that chromosomal *folP*\* genes encoding proteins with the signature Sul motif were independently mobilized into the sul1-3-harboring mobile elements found in sulfonamide-resistant clinical isolates, but they do not address whether the presence of this motif is associated with sulfonamide resistance. To investigate this possibility, we cloned the *folP* gene coding for DHPS in the *Rhodobiaceae P. lavamentivorans* DS-1 (WP_012111048), the *Leptospiraceae L. interrogans* serovar Lai str. 56601 (WP_000444207), the *Rhodobacteraceae R. sphaeroides* 2.4.1 (WP_011337038) and the Chlamydiae *C. trachomatis* D/UW-3/CX (WP_009871981). Following Clinical and Laboratory Standards Institute (CLSI) guidelines [38], we then performed broth microdilution assays to determine the minimal inhibitory concentration (MIC) of sulfamethoxazole. The results shown in Table 1 reveal that both *P. lavamentivorans* and *L. interrogans* chromosomal *folP*\* genes confer resistance to sulfamethoxazole in an *E. coli* strain sensitive to sulfonamides. In contrast, *C. trachomatis folKP* does not confer significant resistance to sulfamethoxazole. Moreover, our results show that complementation with *folP* genes from another Alphaproteobacteria family lacking the Sul motif, the *Rhodobacteraceae*, does not confer resistance. These results reveal that the chromosomal *folP*\* genes that gave rise to *sul* genes are capable of conferring resistance to sulfonamide in *E. coli*. The fact that complementation with *C. trachomatis* and *R. sphaeroides* 2.4.1 *folP*, both lacking the Sul motif, does not confer resistance in the *E. coli* background suggests that sulfonamide resistance in the chromosomal *folP* genes identified here likely originated with protein sequence changes linked to the signature two-amino acid insertion characteristic of mobile *sul* genes and chromosomal *folP*\* genes.

**Table 1.**
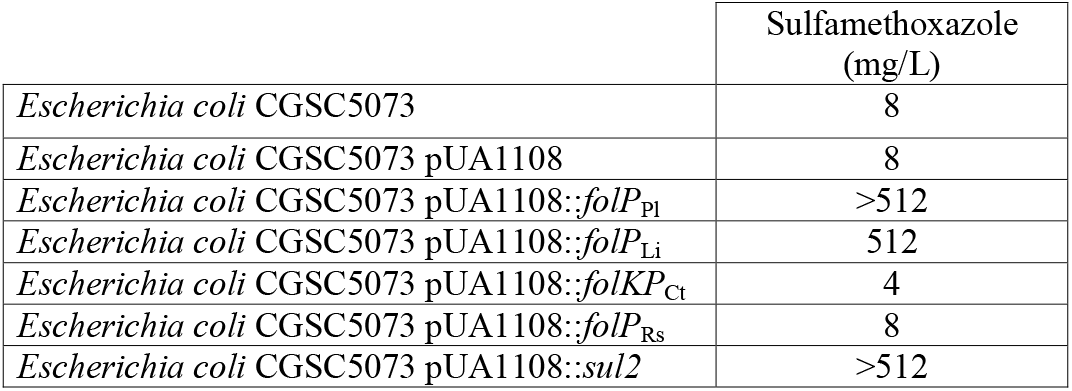
Broth microdilution assays. Minimum inhibitory concentrations (MICs) of sulfamethoxazole in wild-type *Escherichia coli* CGSC5073 carrying different versions of pUA1*108::folP;* Pl, *Parvibaculum lavamentivorans*; Li, *Leptospira interrogans;* Ct, *Chlamydia trachomatis*; Rs, *Rhodobacter sphaeroides*.

## 4 Discussion

### 4.1 Elucidation of the chromosomal origins of *sul1-3* genes

The introduction of sulfonamides in the late 1930’s was soon followed by the emergence of resistance due primarily to mutations in chromosomal *folP* genes [7]. In this context, the most plausible hypothesis for the origin of mobilized *folP* homologs (the *sul* genes) conferring resistance to sulfonamides might appear to involve the uptake by mobile elements of chromosomal *folP* genes that had undergone selection for sulfonamide resistance upon its introduction as a systemic chemotherapeutic agent. Our analysis, however, indicates that the *sul1-3* genes responsible for sulfonamide resistance in clinical isolates did not arise from recently mutated chromosomal *folP* genes. Instead, our results imply that *sul1-3* originated via the independent mobilization of a chromosomal *folP*\* gene that had been horizontally transferred at least once between divergent bacterial clades (Figure 5). This evolutionary scenario is supported by several complementary lines of evidence. The identification of a conserved region incorporating a signature two-amino acid insertion shared by all reported *sul1-3* gene instances and members of the two posited donor families (*Rhodobiaceae* and *Leptospiraceae*) (Figure 1AB) provides strong support for a common origin of these sequences. This result is substantiated by the solidly supported branching of Sul1-3 protein sequences with members of the *Rhodobiaceae* and *Leptospiraceae* families in the reconstructed FolP/Sul molecular phylogeny (Figure 2). Importantly, the trimmed multiple sequence alignment used for FolP/Sul Bayesian phylogenetic inference (Supplementary material 8) does not incorporate the two-amino acid insertion of the Sul motif, indicating that the joint branching of Sul1-3 sequences with chromosomally-encoded *Rhodobiaceae* and *Leptospiraceae* FolP proteins is based on sequence similarity beyond this insertion and its immediate vicinity (Figure 1AB).

**Figure 5.**
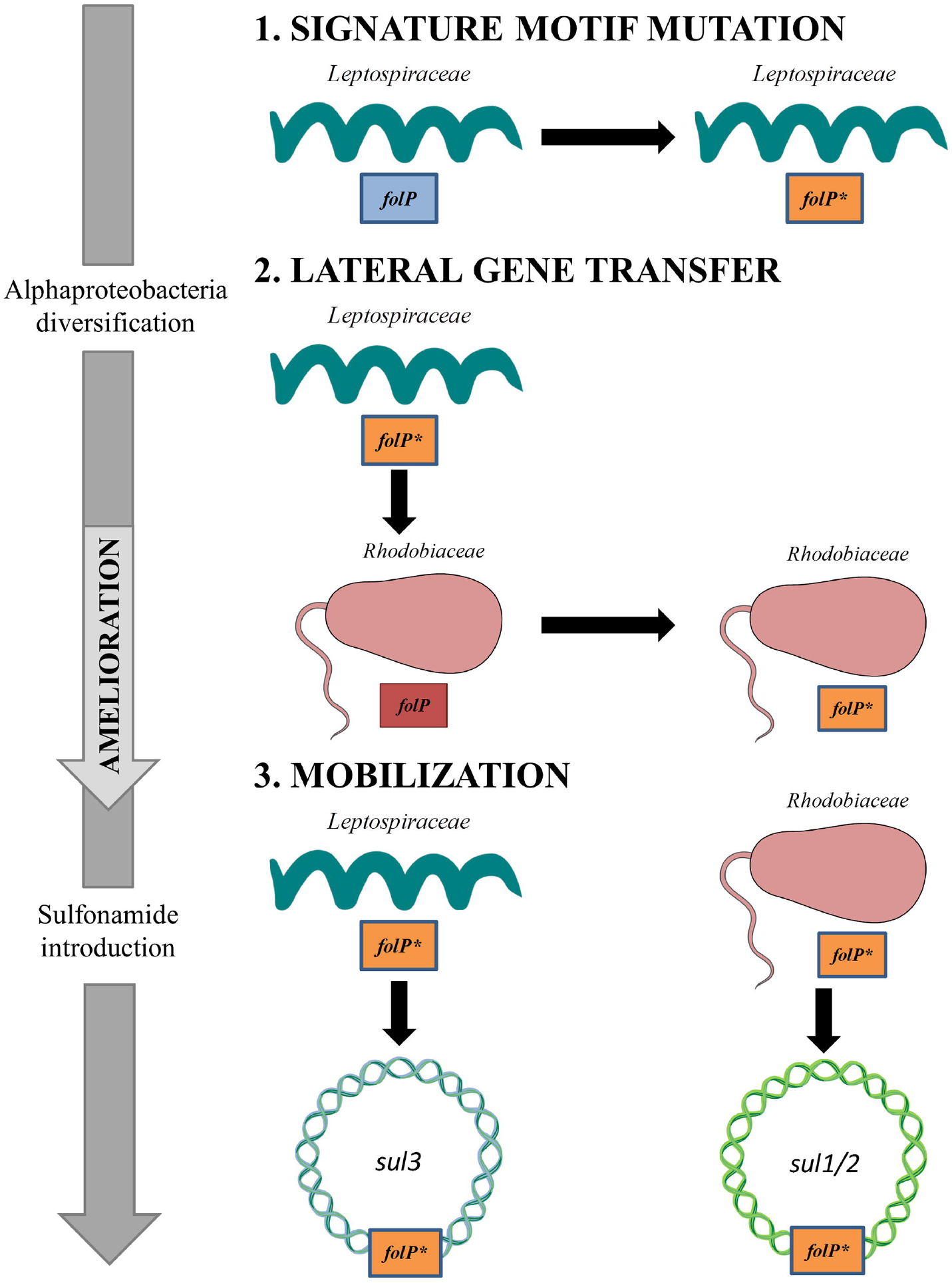
Schematic diagram of the evolutionary process leading to the emergence of *sul1/2*- and *sul3*-harboring mobile genetic elements. (1) A set of mutagenic events in the *Leptospiraceae folP* gene generates the signature motif observed in *folP*\* and *sul* genes. (2) Following the diversification of the Alphaproteobacteria, the *Leptospiraceae folP*\* gene is transferred to the *Rhodobiaceae*. (3) Upon the clinical and agricultural introduction of sulfonamides, folP* genes from the *Leptospiraceae* and the *Rhodobiaceae* are independently mobilized, giving rise to the sul–containing mobile elements reported in clinical isolates. This figure was constructed using some Servier Medical Art templates, which are licensed under a Creative Commons - Attribution Unported License.

The presence of *glmM* gene fragments downstream of *sul2* genes in *sul2* isolates (Supplementary material 1) and the presence of a similar arrangement in the Proteobacteria (Figure 1C) provide an independent means for assessing the origin of *sul2* genes. Phylogenetic inference results for the N-terminal region of GlmM (Figure 3) are consistent with those observed for FolP (Figure 2), and clearly define a last common ancestor between the *Rhodobiaceae* and *sul2*-associated *glmM* genes. Analysis of synonymous and non-synonymous substitutions among *Rhodobiaceae glmM* genes suggests that the *glmM* gene has undergone similar patterns of selection regardless of its association to *folP* genes encoding the signature two-amino acid insertion (Supplementary material 11). Since the *glmM* gene fragment associated to *sul2* genes is likely to be non-functional and subject to genetic drift, the absence of diverging substitution patterns between the N- and C-terminal regions of *Rhodobiaceae glmM* sequences indicates that the *glmM* and *sul2* genes were transferred from the *Rhodobiaceae* to *sul2*-harboring vectors, and not vice versa. Lastly, given that gene loss is much more likely than gain [45], the absence of *glmM* fragments in *sul1* isolates supports in turn the notion that *sul1* derived from *sul2*. This is consistent with the branching pattern observed in the FolP/Sul tree (Figure 2), which defines a scenario of independent mobilization of *sul3* from the *Leptospiraceae* and *sul2* from the *Rhodobiaceae*, with the subsequent uptake of *sul1* by class 1 integrons.

The analysis of *folP* codon %GC content provides further evidence for the evolutionary scenario outlined above (Figure 5). The %GC content of *sul3* genes is very similar to that of *Leptospiraceae folP* sequences, whereas those of *sul2* and *sul1* closely match *Rhodobiaceae folP* genes. Given that more than thirty years elapsed between the introduction of sulfonamides and the detection of *sul*-harboring vectors [7], it is reasonable to assume that *sul* genes were mobilized from chromosomal origins some period of time after the discovery of sulfonamide. Sequence evolution models indicate that, even under fast-evolution scenarios, amelioration from *sul3* to *sul1/2* %GC content (or vice versa) is not feasible in such a short time [32]. In fact, forward simulations suggest that an evolutionary span of at least 476 million years is required to achieve such rates of amelioration. This is congruent with the transfer of *folP*\* from the *Leptospiraceae* to the *Rhodobiaceae* taking place after the inferred diversification of the Alphaproteobacteria into its constituent families some 1,500 million years ago [34]. This timeline is also consistent with the analysis of %GC content, which shows evidence of complete amelioration in *Rhodobiaceae folP*\* genes (Figure 4B). Such an ancestral gene transfer event is also congruent with the lack of canonical telltale signs of lateral gene transfer in either chromosomal background, such as the presence of transposase/integrase genes in the immediate vicinity of *folP*\*, with the substantial diversity of genomic surroundings observed for the *folP* gene in the Alphaproteobacteria (Figure 1C), and with the overlap in habitats between both bacterial families [46,47]. Taken together, these results provide strong support for the hypothesis that the *sul1-3* genes present in clinical isolates were mobilized from chromosomal *Leptospiraceae* and *Rhodobiaceae* backgrounds following the introduction of sulfonamides in the late 1930’s.

### 4.2 Prevalence of sulfonamide resistance in ancestral bacteria

Several independent lines of evidence converge towards an evolutionary scenario in which *sul1-3* genes from clinical isolates derive from ancestral chromosomal mutations in the *folP*\* gene of the *Leptospiraceae* and the *Rhodobiaceae* (Figure 5). This raises several important questions regarding the nature and impact of such chromosomal mutations, the selective pressures underpinning their origin and transfer in ancient bacteria, and their subsequent mobilization into the resistance *sul* genes found in clinical isolates. Minimum inhibitory concentration (MIC) assays confirm that both the *Leptospiraceae* and the *Rhodobiaceae folP*\* genes provide a level of resistance to sulfamethoxazole comparable to that provided by *sul2* gene in an *E. coli* background, whereas complementation with a *Rhodobacteraceae folP* does not confer resistance (Table 1). These data are in agreement with previous reporting of sulfonamide resistance in multiple *L. interrogans* strains [48–50], and suggest that the observed resistance was likely due to mutations in the *Leptospiraceae* chromosomal *folP*\* gene rather than to the presence of plasmid-borne *sul* genes.

In contrast with the *Leptospiraceae* and the *Rhodobiaceae folP*\* genes, the chromosomal *folKP* gene of the Chlamydiae, which encodes a DHPS lacking the Sul motif, does not confer resistance to sulfamethoxazole (Table 1). This is in agreement with abundant reports of sulfonamide susceptibility in several *Chlamydia* species [51–54]. Since the Chlamydiae *folKP* gene is the most closely related chromosomal *folP* gene to the cluster encompassing the *sul* genes and the *Leptospiraceae* and the *Rhodobiaceae folP*\* (Figure 2), the lack of resistance in Chlamydiae *folKP* genes strongly suggests that changes in the region encompassing the Sul motif may be responsible for the observed resistance. This region is located in a connector loop within the N-terminal ‘pole’ of the eight-stranded α/β barrel of DHPS, which is involved in sulfonamide recognition [39,40]. The two-amino acid insertion might hence result in decreased affinity for sulfonamide by locally disrupting folding as has been proposed previously for similar insertions [55].

The emergence and maintenance of a sulfonamide-resistant *folP*\* gene in the *Leptospiraceae* and its subsequent transfer to the *Rhodobiaceae* suggests that it might convey some selective advantage, but the advent of mutations providing significant resistance and their subsequent spread could also have been fortuitous. The appearance of sulfonamide-resistance mutations in chromosomal *folP* genes has been amply documented [7,13], and these were in fact the primary drivers of sulfonamide resistance following the introduction of sulfa drugs [7]. Furthermore, it has been documented that the presence of sulfonamide resistant DHPS does not necessarily impose a fitness cost [56]. Structural studies have suggested that most sulfonamide resistance mutations act by modulating accessibility of sulfonamides to the PABA-binding pocket without hindering PABA binding [40,57].

It is hence conceivable that naturally occurring mutations conferring resistance to sulfonamide might not be selected against in the absence of this chemotherapeutic agent. Subsequent complementary changes to adjust the affinity for PABA of the altered DHPS molecule may have resulted in fixation of the original mutations conferring resistance to sulfonamide [58]. Alternatively, sulfonamide resistance mutations in *folP* may have arisen and persisted in response to naturally occurring sulfonamides produced by competing organisms. Sulfonamides are rare in nature, with only eight known natural sulfonamides reported to date [59]. Of these, only two naturally occurring sulfonamides are aryl sulfonamides, produced in very small amounts by recombinant *Streptomyces* species harboring the complete xiamycin biosynthesis gene cluster [60]. Although these sulfonamides show potent antimicrobial activity, their bulky substitution pattern suggests that their mode of action and molecular target are likely different from synthetic aryl sulfonamides [60].

### 4.3 Mobilization of ancestral resistance reservoirs

The phylogenetic inference and genomic analysis results reported in this work uphold an evolutionary scenario wherein chromosomally-encoded sulfonamide resistant *folP* variants were independently mobilized from *Leptospiraceae* and *Rhodobiaceae* backgrounds following the clinical introduction of synthetic aryl sulfonamides, giving rise to the *sul1/2* and *sul3* genes routinely detected in clinical isolates (Figure 5). The rapid mobilization and dissemination of genes conferring resistance to antibiotic and chemotherapeutic agents upon the clinical or agricultural use of these compounds has been amply documented [4,61]. Mobilization and spread may be mediated by plasmids encoding transposons and integrons, as well as integrative and conjugative elements, mobile pathogenicity islands and bacteriophages, but the common tenet is that sustained exposure of bacterial populations to antibiotics or chemotherapeutic agents induces a strong selective pressure to elicit the mobilization of resistance determinants [61].

Together with penicillin and tetracycline, sulfonamides have been the antibacterial agents most frequently used at sub-therapeutic levels in livestock production [62], and it has been reported that sulfonamides have higher mobility, low removal efficiency and deeper environmental penetration than most other antibacterial agents [63]. The widespread and intensive use of sulfonamides in agriculture, aquaculture and animal husbandry since the mid 1960’s, and their persistence in soil, sediments and subterranean aquatic communities where *Leptospiraceae* and *Rhodobiaceae* abound, provides an ample window of opportunity for the mobilization of chromosomally-encoded *folP*\* genes within these bacterial communities and the subsequent transfer of these mobile resistance determinants to other bacteria.

Recent mobilization from a Chloroflexi chromosomal *folP* background has been postulated as the likely origin of the *sul4* gene [22], and this result is in agreement with the phylogenetic analysis reported here (Figure 2). In the case of the chromosomal *folP*\* identified here and their mobilization into *sul*-harboring resistance vectors, several sources of evidence provide additional support for the mobilization of chromosomal *folP* genes. For instance, phylogenetic evidence (Figure 2) indicates that the *Rhodobiaceae folP*\* was incorporated at some point by the Actinobacterium *Amycolatopsis*, which harbors three *folP* orthologs (Supplementary material 12). Similarly, a plasmid broadly distributed among *Azospirillum* plasmids (e.g. AP010951, FQ311873), a member of the *Rhodospirillaceae* Alphaproteobacteria family, contains a *folP* gene flanked by genes coding for a flagellar export pore protein (FlhB) and the full length phosphoglucosamine mutase (GlmM) (Supplementary material 12). This *folP* does not contain the signature two-amino acid insertion, indicating that its mobilization occurred independently of those leading to *sul1/2* genes.

More significantly, a partial genomic sequence from a *Pseudomonas aeruginosa* isolate (LLMY01000073.1) harbors a *folP*\* gene with high sequence and genetic neighborhood similarity to the *Rhodobiaceae P. lavamentivorans* DS-1 [64]. The genes immediately upstream and downstream of this *P. aeruginosa folP*\*, which contains the *sul* motif, encode a TetR family regulator and a partial phosphoglucosamine mutase (GlmM) protein (Supplementary material 12). These three genes are flanked by IS91 and ISL3 family transposases. Importantly, the IS91 transposase contains similar sequence motifs and shares termini identity with ISCR elements, which are present in both *sul1* and sul2-harboring plasmids [65,66]. It is hence highly likely that the *P. aeruginosa folP* represents an intermediate step in the original mobilization of *sul1/2* from a *Rhodobiaceae* background.

Metagenomics analysis and prospective studies of preserved ancient environments, such as permafrost and remote cave habitats, have largely displaced the notion that antibiotic resistance emerges in response to anthropogenic antibiotic use [67–70]. These studies have conclusively shown that antibiotic resistance predates the use of antibiotics by humans, and that it is widely distributed across the bacterial pangenome. In a few isolated cases, resistance determinants for synthetic chemotherapeutic agents that predate or have rapidly arisen upon human use has been documented, but their existence can be attributed to cross-resistance to naturally-occurring antibiotics (e.g. microcin B17 for quinolones [6], sisomicin for amikacin [69]). The identification in this work of ancient chromosomal mutations in *folP* conferring resistance to sulfonamide as the likely origins of the *sul1-3* genes present in sulfonamide-resistant clinical isolates puts forward an alternative scenario. Given the absence of known naturally occurring aryl sulfonamides targeting DHPS, our results suggest that resistance to novel synthetic chemotherapeutic agents may be already available in the vast microbial pangenome, and that its mobilization and global dissemination can take place in a very short amount of time upon the clinical introduction of novel chemotherapeutic compounds.

## 7 Author Contributions

MSO and IE performed the *in silico* analyses. MSO and IE developed scripts for genomic analyses and ran phylogenetic inference methods. MSO and PC performed the *in vitro* analyses. All authors discussed the findings and interpreted the results. IE and JB conceived the experiment and coordinated the research. IE and MSO drafted the manuscript.

## 9 Acknowledgments

The authors wish to thank Joan Ruiz and Susana Escribano for their technical support during some of the experimental procedures, as well as Júlia López for her continued support.

## 10 Supplementary Material

The Supplementary Material for this article can be found online at:

## 11 Data Availability Statement

The datasets used in this study can all be freely accessed at the NCBI GenBank/RefSeq databases (https://www.ncbi.nlm.nih.gov/). All scripts used for analysis can be obtained at the GitHub ErillLab repository (https://github.com/ErillLab/).

## 14 Supplementary material

**Supplementary material 1** – List of accession numbers for chromosomal and plasmid sequences containing FolP/Sul/GlmM-encoding genes used in this work. The sequence accession number, the species name and the corresponding FolP, Sul and GlmM protein accessions are provided in different columns.

**Supplementary material 2** – List of oligonucleotides used in this work.

**Supplementary material 3 –** Multiple sequence alignment including *sul* genes at most 99% similar to those reported in the literature and one representative chromosomal *folP* gene from bacterial phyla with complete genomes available in RefSeq.

**Supplementary material 4** – PROSITE-formatted pattern of the region containing the identified two-amino acid insertion in sul-encoded proteins used to seed the PHI-BLAST search.

**Supplementary material 5** – Detail of the multiple sequence alignment region containing the two-amino acid signature motif including Sul sequences and FolP sequences from members of the *Rhodobiaceae*, the *Leptospiraceae* and the Chlamydiae.

**Supplementary material 6** – List of accession numbers (Nucleotide and Protein) for the *sul1, sul2* and *sul3* genes used in the tBLASTX search of *folP* genetic surroundings.

**Supplementary material 7** – Unrooted Neighbor-Joining tree of Sul/FolP homologs. Branch support values are provided as the percent of bootstrap replicates in which the branching was observed. Support values are only shown for branches with at least 80% support. The cluster of Actinobacteria duplicated *folP* gene products that were removed from further analysis is indicated by the shaded ellipse.

**Supplementary material 8** – Multiple sequence alignment (FASTA format) of FolP/Sul sequences used for phylogenetic inference after trimming with GBLOCKS.

**Supplementary material 9** – Sequences of *folP* genes used in the *folP* sequence analysis (FASTA format).

**Supplementary material 10** – Values of overall and codon-position %GC content across all protein coding genes in a genome, and for *folP* genes, on genomes of all bacterial orders with complete genome assemblies, of each bacterial family for the Proteobacteria and all available complete genomes for clades of interest (*Rhodobiaceae*, Spirochaetes and Chlamydiae).

**Supplementary material 11** – Synonymous and non-synonymous mutation patterns in pair-wise alignments between the N- and C-terminal regions of *Rhodobiaceae glmM* genes.

**Supplementary material 12** – Schematic representation of the genetic environment of *sul2* genes, similar arrangements in chromosomally-encoded *folP* genes of the Gammaproteobacteria, Betaproteobacteria and Alphaproteobacteria, and arrangements in other putative mobilization instances of the *folP* gene. Arrow boxes indicate coding regions. When available, gene names or NOG identifiers are provided. Boxes for *folP* genes containing the two-amino acid insertion are designated as *folP*\*.

